# Virtual partition digital PCR for high precision chromosomal counting applications

**DOI:** 10.1101/2021.04.29.441975

**Authors:** Lucien Jacky, Dominic Yurk, John Alvarado, Bryan Leatham, Jerrod Schwartz, Chris MacDonald, Aditya Rajagopal

## Abstract

Digital PCR (dPCR) is the gold standard analytical platform for rapid high precision quantification of genomic fragments. However, current dPCR assays are generally limited to monitoring 1-2 analytes per sample, thereby limiting the platform’s ability to address some clinical applications that require the simultaneous monitoring of 20 – 50 analytes per sample. Here we present Virtual Partition dPCR (VPdPCR), a novel analysis methodology enabling the detection of 10 or more target regions per color channel using conventional dPCR hardware and workflow. Furthermore, VPdPCR enables dPCR instruments to overcome upper quantitation limits caused by partitioning error. While traditional dPCR analysis establishes a single threshold to separate negative and positive partitions, VPdPCR establishes multiple thresholds to identify the number of unique targets present in each positive droplet based on fluorescent intensity. Each physical partition is then divided into a series of virtual partitions, and the resulting increase in partition count substantially decreases partitioning error. We present both a theoretical analysis of the advantages of VPdPCR and an experimental demonstration in the form of a 20-plex assay for non-invasive fetal aneuploidy testing. This demonstration assay – tested on 432 samples contrived from sheared cell-line DNA at multiple input concentrations and simulated fractions of euploid or trisomy-21 “fetal” DNA – is analyzed using both traditional dPCR thresholding and VPdPCR. VPdPCR analysis significantly lowers variance of chromosome ratio across replicates and increases the accuracy of trisomy identification when compared to traditional dPCR, yielding *>*98% single-well sensitivity and specificity. VPdPCR has substantial promise for increasing the utility of dPCR in applications requiring ultra-high-precision quantitation.

## Introduction

In many clinical diagnostic applications, it is essential to not only detect the presence of a nucleic acid target, but to also measure its concentration. This is commonly done with the quantitative polymerase chain reaction (qPCR), which calculates concentration based on the number of PCR cycles needed for a sample to reach a certain signal threshold. This method benefits from widespread instrument deployment and a simple workflow, but requires running standards for calibration purposes and is only effective at establishing target concentration within a factor of 2 [1–3].

An alternative approach to target quantitation is digital PCR (dPCR). dPCR divides a sample into tens of thousands of individual partitions prior to amplification. After amplification, the partitions that generate an amplification signal are counted to produce measurements of target concentration. When compared to qPCR, dPCR improves both precision and low-copy sensitivity, while providing absolute quantitation without the need for calibrating standards [1–4]. Due to its advantages, dPCR has shown increasing utility in a number of diagnostic applications, including absolute quantification of viral load, analysis of circulating DNA, gene and microRNA expression, and analysis of gene copy number variation [5–9].

The most fundamental limit to the precision of dPCR quantitation is Sampling Variance; if the true mean number of target copies across many sample replicates is *N*, the standard deviation of the number of copies truly present in each replicate will be 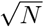 [10] (Figure 1a). For applications that require high precision at low input copy numbers, this level of variance can be unacceptably high. The number of effective sample copies can be increased by multiplexing, i.e. designing multiple assays specific to different regions of the target. However, as the number of effective sample copies passes the number of physical partitions the partitions become oversaturated, leading to high levels of Partitioning Variance [10]. This tradeoff, illustrated in Figure 1b, has limited the usefulness of multiplexing for decreasing Sampling Variance in single-well dPCR assays [5].

**Fig 1.**
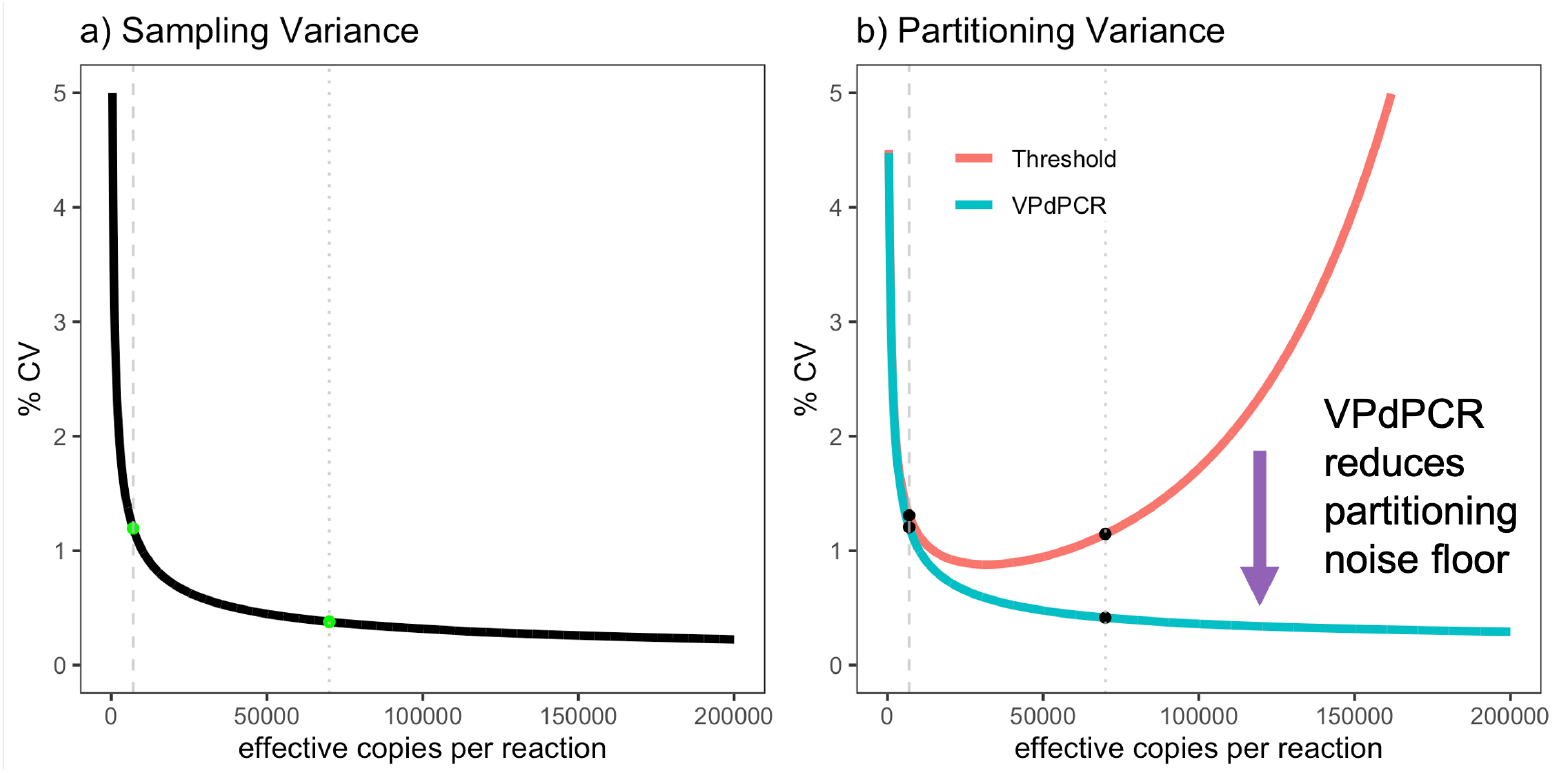
Digital PCR mathematical noise floor. The coeffecient of variation (CV) of calculated copies per reaction across a range of expected copies in sample, assuming a digital PCR reaction with 20,000 physical partitions. The dashed grey line represents 7,000 copies per reaction with only one assay per target, while the dotted grey line represents 70,000 effective copies corresponding to assays for 10 unique target regions on each of 7,000 target copies. a) Sampling Variation: The standard deviation due to sampling error is defined as the square-root of the mean expected copies in a sample. This error is independent of the analysis method. Increasing the number of assays per target effectively increases the expected number of copies and decreases the Sampling Variance. b) Partitioning Variation: The standard deviation due to partitioning error is dependent on the number of partitions a sample is divided into. The VPdPCR assay (blue) described in this paper increases the effective number of partitions 10-fold, significantly reducing the mathematical noise floor due to partitioning error when compared with the tradition threshold method (red).

The statistical analysis underlying the red line in Figure 1b assumes traditional dPCR analysis, which uses a single amplitude threshold to classify each partition as either negative for all targets or positive for at least one target. One way around this Partitioning Variance limitation is to apply enhanced assay design to generate multiple distinct signal clusters corresponding to different combinations of target regions in the partitions. Historically this process has proven difficult, limiting such multiplexing applications to 2 or 3 targets per optical color channel [11–15]. However, this difficulty can be overcome by applying High Definition PCR (HDPCR^™^), a recent innovation in qPCR technology, which has been used to expand the multiplexing capacity of qPCR instruments [16]. In this paper we describe Virtual Partition dPCR (VPdPCR), a method which leverages HDPCR and a novel analysis technique to enable significantly higher levels of multiplexing on existing dPCR instruments using standard TaqMan chemistries. We present both a theoretical analysis of the advantages of VPdPCR and an experimental demonstration of its capabilities in the form of a single-well, 20-plex assay for detection of fetal chromosomal aneuploidies.

### Virtual Partition dPCR

VPdPCR combines multiple TaqMan^®^ assays that are designed to detect multiple distinct regions of the desired target, in this case a particular chromosome. The TaqMan probes for all target regions on a given chromosome are labeled with the same fluorophore and quencher pair, and they are all titrated to generate the same fluorescence intensity if the target region is present. These probes generate signals that add linearly in combination; if a single target region generates fluorescence intensity *I* when it is present in a partition, a partition with *n* distinct target regions will have fluorescence intensity *n* I*. The number of target regions presents in each partition can thus be inferred solely from the signal intensity measurement of the partition.

In traditional dPCR analysis, a single signal intensity threshold is drawn to separate partitions positive for 1 or more target regions from those partitions negative for all target regions. VPdPCR changes the readout of a digital partition from a 2 state system to a *T* + 1-state system by drawing *T* different intensity thresholds, where *T* is the number of target regions and 1 is added for the negative state (Figure 2). Each partition is divided into *T* “virtual partitions”, enabling higher allowable target concentrations without oversaturation by increasing the number of effective partitions by a factor of T. Creating these virtual partitions significantly reduces the Partitioning Variance which usually occurs as the occupancy of the physical partitions approaches 100% (Figure 1b, blue curve).

**Fig 2.**
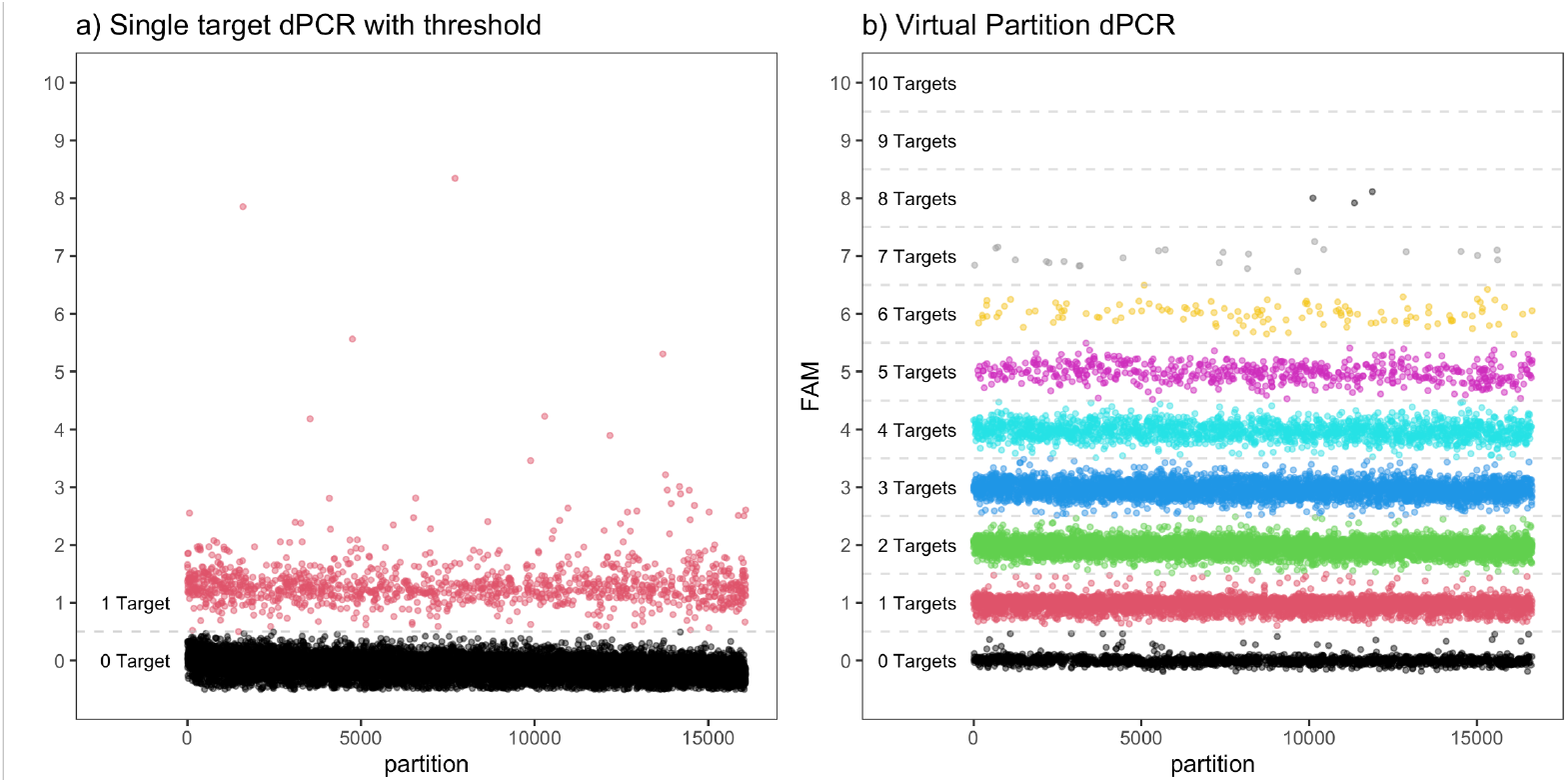
Evolving Multiplex Digital PCR. a) Traditional singleplex dPCR is a robust method using a simple threshold but is limited by significant variation at low input and high input due to sampling and Partitioning Variance respectively. b) By combining VPdPCR with HDPCR we’re able to divide each partition into many bins to determine the copies per virtual partition. This substantially reduces Partitioning Variance by expanding the number of effective partitions.

This manuscript presents a demonstration assay capable of detecting 10 unique target regions per channel, with one channel devoted to targets from chromosome 18 and the other dedicated to targets from chromosome 21 (Figure 3). By increasing the effective target concentration, VPdPCR drives down Sampling Variance while avoiding high Partitioning Variance caused by oversaturation (Figure 1). This allows for consistent detection of very small differences in abundance ratio between the chromosomes. To demonstrate the utility of the technique, we apply it to the problem of differentiating simulated cell-free DNA (cfDNA) samples with and without small fractions of trisomy 21 DNA.

**Fig 3.**
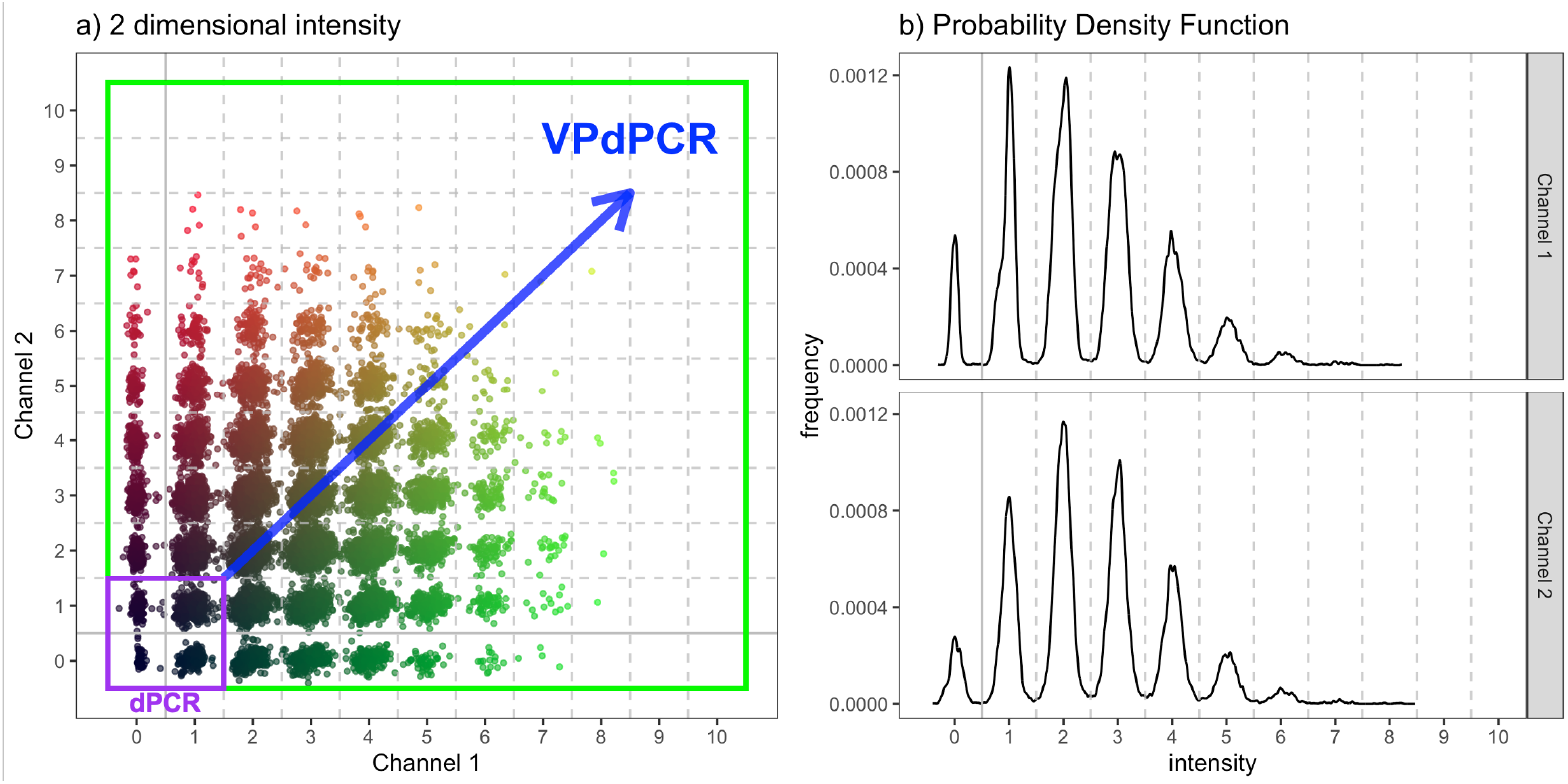
Virtual Partition dPCR data. a) Two dimensional plot of experimental VPdPCR data with 7000 haploid genomic copies in 20,000 physical partitions. The assay was designed with 10 independent target regions per chromosome, with chr18 target regions in channel 1 and chr21 target regions in channel 2. Thanks to HDPCR, distinct point clusters can be discerned containing droplets with different numbers of target regions. The color of the individual points represents the intensity of channel 1 and channel 2 as the Green and Red component of RGB values respectively. The purple box encompasses the partitions that can be analyzed with standard dPCR thresholding, while the green box encompasses the additional virtual partitions which can be interrogated in a 10 target per channel VPdPCR assay. b) Probability density function (PDF) plots for channel 1 and channel 2 of the experimental data from the same reaction. Each channel’s PDF is analyzed independently to calculate copies of each chromosome. In both plots the solid grey line represents the traditional threshold between positive and negative physical partitions, while the dashed grey lines separate the virtual partitions differentiated by signal intensity.

### Non-Invasive Fetal Aneuploidy Testing

Screening for fetal aneuploidy in expectant mothers is one of the most common forms of prenatal diagnostics in the world [17], and is traditionally performed using methods such as chorionic villus sampling or amniocentesis. While these tests still represent a gold standard for accuracy, their invasive nature limits their application to high-risk populations [18]. The discovery of fetal cell-free DNA (cfDNA) circulating in maternal blood [19] opened the door to non-invasive prenatal testing (NIPT) for fetal aneuploidy by counting chromosomal copies; if a fetus is triploid for a given chromosome, the number of copies of that chromosome in fetal cfDNA should be 50% higher than all other copy numbers. Modern fetal aneuploidy tests are often performed via next-generation sequencing or microarray tests which require expensive equipment and consumables as well as complicated multi-day workflows, largely relegating them to centralized laboratories and driving up costs.

dPCR provides lower cost, lower complexity, and higher throughput when compared to NGS or microarray tests, making it a desirable modality for fetal aneuploidy screening. However, no single well dPCR-based assay for NIPT has come to market [20]. The primary reason is limited precision; for a euploid mother and an aneuploid fetus with a trisomy only the fetal portion of the cfDNA will show an excess in chromosomal copies, and the fraction of fetal cfDNA derived from a maternal blood or plasma sample can be as low as 4% [21]. This results in only a 2% excess of the fetal trisomy chromosome amongst the whole cfDNA sample, and standard single target dPCR assays are unable to distinguish maternal from fetal DNA. As a result, attempting to consistently measure the excess of the fetal trisomy chromosome using single-plex dPCR proves especially difficult due to Sampling Variance.

Multiple groups have attempted to use multiplexing to bypass this Sampling Variance problem [22–24]. However, in order to avoid oversaturation, these assays must increase the number of effective partitions by either splitting each sample across 8 or more wells [22, 24] or using specialized (and now-discontinued) platforms that can generate millions of partitions per sample [23]. Both of these approaches increase cost and decrease throughput. We instead apply VPdPCR to increase the number of effective partitions, thereby substantially improving accuracy of quantitation at higher target concentrations than was previously possible in a single-well assay. While the presented results are only intended as a proof of concept, they establish the power of VPdPCR as a potential foundation for future ultra-high-precision quantitative assays.

### Glossary

- Target: the whole nucleic acid molecule of which the concentration is being interrogated. Examples: a whole genome, an individual chromosome, a particular RNA transcript.
- Target Region: a sub-sequence within the complete Target sequence which is detected by a unique assay. Example: the template region of a Target for a PCR detection assay.
- Partition: one of many independent physically separate PCR reactions into which a sample is equally divided in dPCR. Examples: A individual droplet or microfluidic well.
- Virtual partition: Expanded partitions derived from the signal amplitude when multiple Target Region assays are leveled to produce the same signal intensity. Example: in an assay with 10 Target Regions to interrogate a single Target, every physical partition is divided into 10 virtual partitions using the VPdPCR method.
- Classification: Analyzing a partition and determining how many positive targets it contains based on its signal amplitude. Examples: identifying a partition as positive or negative in traditional dPCR, or placing it into a bin level 0 through T in VPdPCR.
- Positive Partition Count: the total number of positive partitions identified after classification. Examples: The sum of positive physical partitions in traditional dPCR or the sum of positive virtual partitions in VPdPCR.
- Target Region Copies: the computed total number of target regions present in a sample after obtaining a Positive Partition Count and applying Poisson statistics. Example: Chr18 Copies is the imputed total number of target regions from chromosome 18 present in the original sample.
- Sampling Variance: The variation the number of targets which actually end up in a reaction due the sub-sampling of a larger population. This has larger effect on low concentration sample accuracy due to the standard deviation being 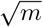 where m is the expected number of targets [25].
- Partitioning Variance: The variance attributed to the distribution of targets between the partitions. At high concentrations the number of empty partitions the standard deviation 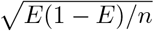 where E is the proportion of negative partitions and n is the number of partitions [10].

## Results and Discussion

We contrived 432 cell line-based DNA samples, each consisting of a mixture of “maternal” euploid DNA from a wild type cell line and simulated “fetal” DNA from either a trisomy 21 cell line or a different euploid cell line. Half of the samples contained 7000 total copies of chr18 and half of the samples contained 3500 copies – both of which are representative of DNA concentrations in a typical cell-free DNA extraction from expectant women [26]. Within each of these sets, 36 samples each had simulated aneuploid trisomy 21 fetal fractions of 0%, 5%, 10%, and 20%, and 36 samples each had simulated euploid fetal fractions of 5% and 100%. Two of the 20% simulated aneuploid fetal fractions wells at 7000 input copies had failures in the droplet reader and results were discarded. Summary results are shown below in Table 1 with all euploid samples condensed into one row, and these euploid samples are broken out by cell line composition in Table 2. An unexpected result is that even in the purely euploid samples the chr21 counts are 4.9% higher than chr18 counts on average. There are multiple possible explanations for this, including off-target amplifications, a duplicated target region on chr21, or chr18 target regions being more susceptible to shearing during sample preparation. Fortunately, this excess is consistent across all euploid sample compositions and the difference in ratios between this baseline and other experimental conditions (ΔRatio in the table) scales as expected with aneuploid fraction, indicating that this is a consistent offset across all samples. Thus, when classifying samples to identify fetal trisomy we treat 1.049 as the baseline euploid chromosome ratio.

**Table 1.** Contrived Trisomy Pregnancy Results.

| Sim. Aneuploid Fraction | Sample Size | VPdPCR Analysis |  |  |  |  |  | Threshold Analysis |  |
| --- | --- | --- | --- | --- | --- | --- | --- | --- | --- |
| | | $\bar{\lambda}_{\text{Chr18}}$ | $\bar{\lambda}_{\text{Chr21}}$ | Ratio* | ΔRatio | $\sigma_{\text{Ratio}}$ | Accuracy | $\sigma_{\text{Ratio}}$ | Accuracy |
| 7k Haploid Genomic Copies per Well |  |  |  |  |  |  |  |  |  |
| 0%** | 108 | 0.2666 | 0.2794 | 1.048 | -0.1% | 0.86% | 98.1% | 1.46% | 85.1% |
| 5% | 36 | 0.2658 | 0.2864 | 1.078 | 2.9% | 0.83% | 97.2% | 1.72% | 77.8% |
| 10% | 36 | 0.2641 | 0.2912 | 1.103 | 5.4% | 0.67% | 100% | 1.58% | 100% |
| 20% | 34 | 0.2632 | 0.3037 | 1.154 | 10.5% | 0.86% | 100% | 1.51% | 100% |
| 3.5k Haploid Genomic Copies per Well |  |  |  |  |  |  |  |  |  |
| 0%** | 108 | 0.1325 | 0.1390 | 1.049 | 0.0% | 1.02% | 98.1% | 1.28% | 87.0% |
| 5% | 36 | 0.1330 | 0.1435 | 1.080 | 3.1% | 0.95% | 88.9% | 1.39% | 86.1% |
| 10% | 36 | 0.1337 | 0.1475 | 1.103 | 5.4% | 0.80% | 100% | 1.33% | 100% |
| 20% | 36 | 0.1327 | 0.1532 | 1.154 | 10.5% | 1.10% | 100% | 1.48% | 100% |

\* $\lambda_{\text{Chr21}}/\lambda_{\text{Chr18}}$
\*\*Includes the three euploid only Cell line DNA mixtures from the Table 2: Contrived Euploid Pregnancy Results.

**Table 2.**
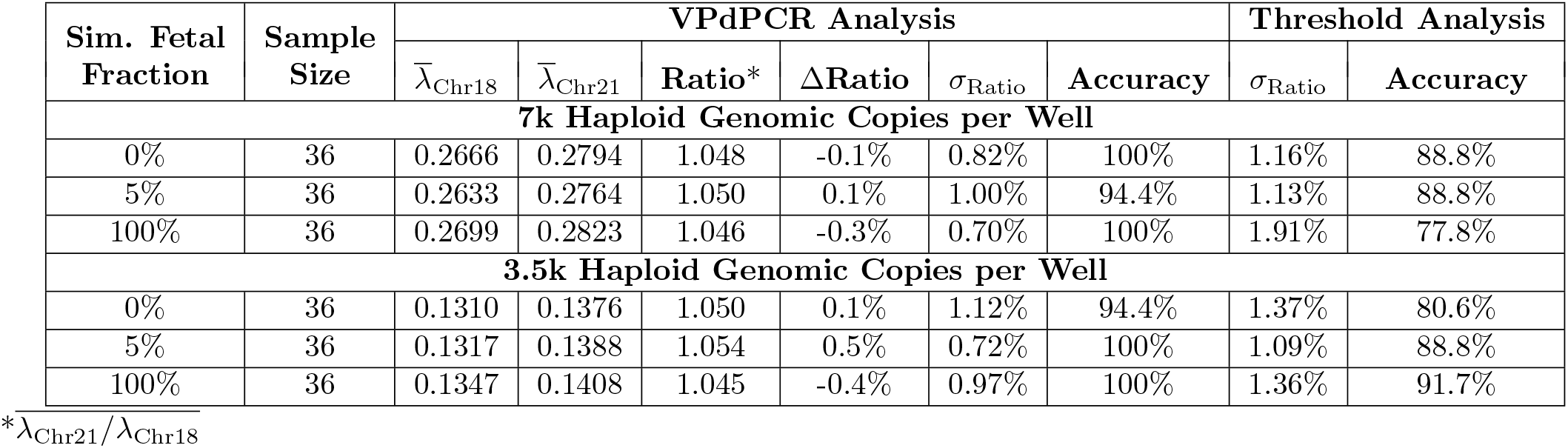
Contrived Euploid Pregnancy Results.

Receiver operating characteristic (ROC) analysis was performed on the complete data set to identify the optimal threshold to separate trisomy 21 spiked samples from the euploid samples using the ratio of Chr21/Chr18 as the predictor. The calculations were performed using the R software package pROC [27, 28]. The optimal threshold was determined to be 1.0672 (Figure 4).

**Fig 4.**
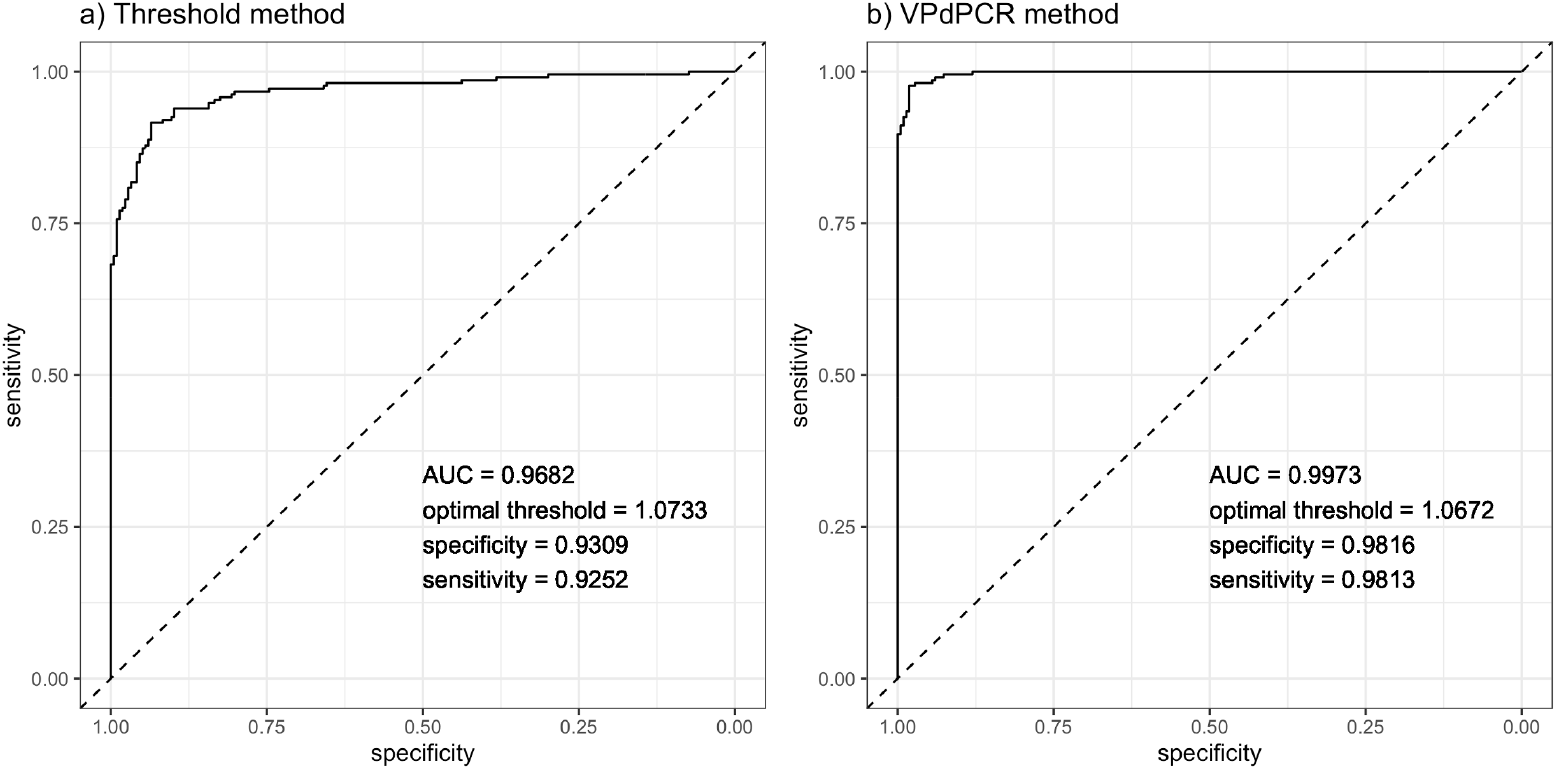
Receiver operating characteristic curve. ROC curves generated using a) the traditional threshold method and b) VPdPCR method. Both ROC curves use the the complete data set including all fetal fractions and both starting concentrations to determine the optimal threshold to separate the pure euploid from trisomy 21 spiked samples.

In the tables above *σ*_Ratio_, or the standard deviation of calculated chromosome ratio across all replicates, is the most important metric for determining the level of quantitative accuracy the assay has achieved. A lower *σ*_Ratio_ indicates that the assay is more able to precisely identify the true ratio of chromosomes present in the sample, thereby increasing its accuracy in high-precision applications like fetal trisomy testing.

Tables 1 and 2 compare *σ*_Ratio_ when samples were analyzed using VPdPCR (Equation 2) versus the traditional method (Equation 1). The traditional analysis was conducted using sample-specific positive/negative amplitude thresholds for each well by taking the midpoint of the fitted 0-target and 1-target peaks in each channel. Even with this optimized thresholding, the VPdPCR analysis consistently achieved lower *σ*_Ratio_ on every set of replicates when compared to traditional analysis. This difference was most pronounced for 7k input samples, where VPdPCR cut *σ*_Ratio_ by more than a factor of 2 in some cases. For 3.5k input samples we expect VPdPCR to provide less of an advantage, as partitioning error is less pronounced at lower input concentrations due to less oversaturation. This theory is reflected in the results, which show a smaller but still consistent improvement from applying VPdPCR in these samples. The differences in ratio distributions from the two analyses are shown visually in Figure 5.

**Fig 5.**
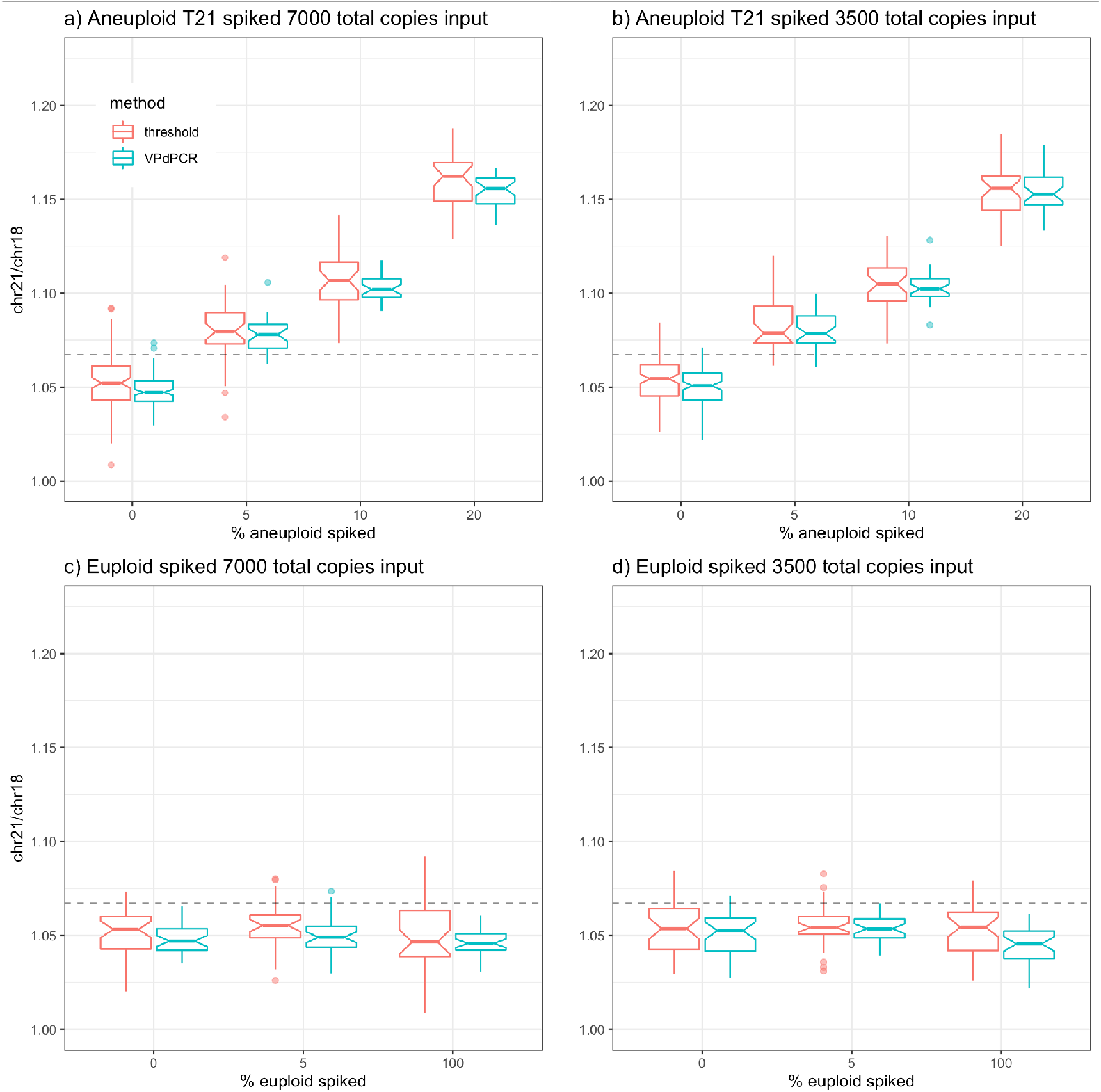
Ratio of Chromosome 21 to Chromosome 18. The ratio Chr21 to Chr18 across a range of euploid and trisomy 21 DNA spiked into euploid background DNA using both the traditional threshold (red) and virtual partition (blue) analysis methods. a) Samples spiked with a T21 cell-line DNA into a euploid background with 7000 haploid genomic equivalents of chromosome 18 per reaction. b) Samples spiked with a T21 cell-line DNA into a euploid background with 3500 haploid genomic equivalents of chromosome 18 per reaction. c) Samples spiked with a euploid cell-line DNA into a different euploid cell-line background with 7000 haploid genomic equivalents of chromosome 18 per reaction. d) Samples spiked with a euploid cell-line DNA into a different euploid cell-line background with 3500 haploid genomic equivalents of chromosome 18 per reaction. In the box and whiskers plots the center line corresponds to the median, the lower and upper boxes represent the first and third quartiles respectively, and the whiskers extend from the boxes to the smallest and largest values no further from the median than 1.5 times the inter-quartile range. Outlier data beyond the whiskers is represented by individual points and the notch within the boxes approximates the 95% confidence interval of the median. The dashed line is the optimal threshold of 1.0672 as determined by ROC analysis for all samples using the VPdPCR method. Plots were generated with the R software package ggplot2 [29].

Table 1 show the accuracy of our assay when applying the ROC-optimized threshold to separate euploid from aneuploid samples. 0% simulated fetal fractions samples were called correct if the computed chr21/chr18 ratio fell below the ROC-threshold, and all other samples were called correct of their ratio fell above the threshold. As shown in Figure 5 the ratio distributions for 0% and 5% simulated fetal fractions overlap significantly less when VPdPCR analysis is applied, and this is reflected in the accuracy results; VPdPCR consistently classifies samples with higher accuracy than traditional analysis does. It is worth emphasizing that this data represent a proof of concept for the VPdPCR technique and are not intended to demonstrate clinical viability. This proof of concept demonstrates that VPdPCR has substantial promise for increasing the utility of digital PCR in applications like fetal trisomy screening requiring ultra-high-precision quantitation.

## Materials and methods

### Novel dPCR Analysis: Multi-Gaussian Fitting

In dPCR analysis the goal is to determine the number of copies per partition of each target, denoted by *λ*. For multi-channel assays, if we assume targets are independently distributed we can treat each channel independently and thereby compute a separate *λ* for each channel; this approach is taken for all presented analyses (Figure 3b). The distribution of each target amongst all partitions is dictated by Poisson statistics, which specify that the probability of a partition being negative for a target with concentration *λ* is simply *p*_*neg*_ = *e*^−*λ*^. In traditional single-target dPCR analysis a single amplitude threshold is drawn to separate positive from negative partitions, and target concentration (in copies per partition) is calculated as *λ* = − ln (*p*_*neg*_) (where *p*_*neg*_ is the fraction of partitions below the threshold). If *T* targets are present at identical concentrations these equations become

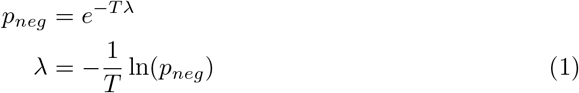

In real experiments, the precision with which we can determine *p*_*neg*_ is limited by the number of negative partitions, with fewer negative partitions leading to higher variance.

This is what leads to the oversaturation effect described earlier. For example, if *T* = 10 and *λ* = 1 we get *p*_*neg*_*≈* 5 *×*10^−5^, which corresponds to only a single negative partition on average in a 20,000 partition system. This leads to extremely high errors with traditional analysis, necessitating new analysis techniques in such a regime.

To perform more accurate analysis, we need to identify how many of the *T* targets are present in each partition rather than merely determining whether or not the partition is negative for all targets. This can be done by dividing up the amplitude range of our partitions into bins with indices *t* = 0, 1, …, *T*. Once bin boundaries are determined, the number of targets present in each partition can be counted by simply determining which boundaries its amplitude falls in between. In principle, on a perfectly consistent system one could run calibration wells and manually draw boundaries between all peaks, then apply those boundaries to sample data. However, on real data, peak locations vary from sample to sample due to a combination of instrument and pipetting variance, leading to poor performance with fixed bin boundaries.

We instead use a more robust method of multi-peak fitting which takes advantage of two observed properties of the system: 1) each peak in the probability density function (PDF) of partition amplitudes can be well approximated by a Gaussian function 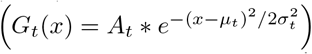 and 2) peak amplitudes add linearly, with equal spacing between each subsequent pair of peaks. The fit is based on 5 free parameters: Target Region Copies per droplet (*λ*, assumed to be the same for all target regions on a chromosome); centers of the 0-target and 1-target bins (*µ*_0_ and *µ*_1_); and widths of the 0-target and 1-target bins (*σ*_0_ and *σ*_1_) The linearity of the system allows us to determine the center and width of all subsequent peaks:

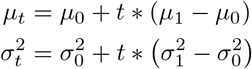

We can also determine the heights of all peaks based off of 1) Poisson statistics that dictate the probability *P* (*t*) of a partition containing *t* target regions and 2) the fact that the area under a Gaussian curve is equal to 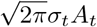

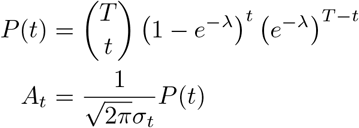

Once we have calculated *µ, σ*, and *A* for every peak we construct a full predicted PDF by adding all of the Gaussian functions together (Figure 6a). The optimal set of (*λ, µ*_0_, *µ*_1_, *σ*_0_, *σ*_1_) is determined to be the one which minimizes the RMS error between the full predicted PDF and the observed PDF. This fit is then used to determine *n*(*t*), the total number of partitions in each bin. Rather than assigning each one of the *N* total partitions to a single bin, we divide it between bins based on the relative magnitude of each bin’s Gaussian at the partition’s amplitude (*x*_*i*_), improving classification accuracy for higher-order bins in which the tails of neighboring Gaussians start to blend into each other:

**Fig 6.**
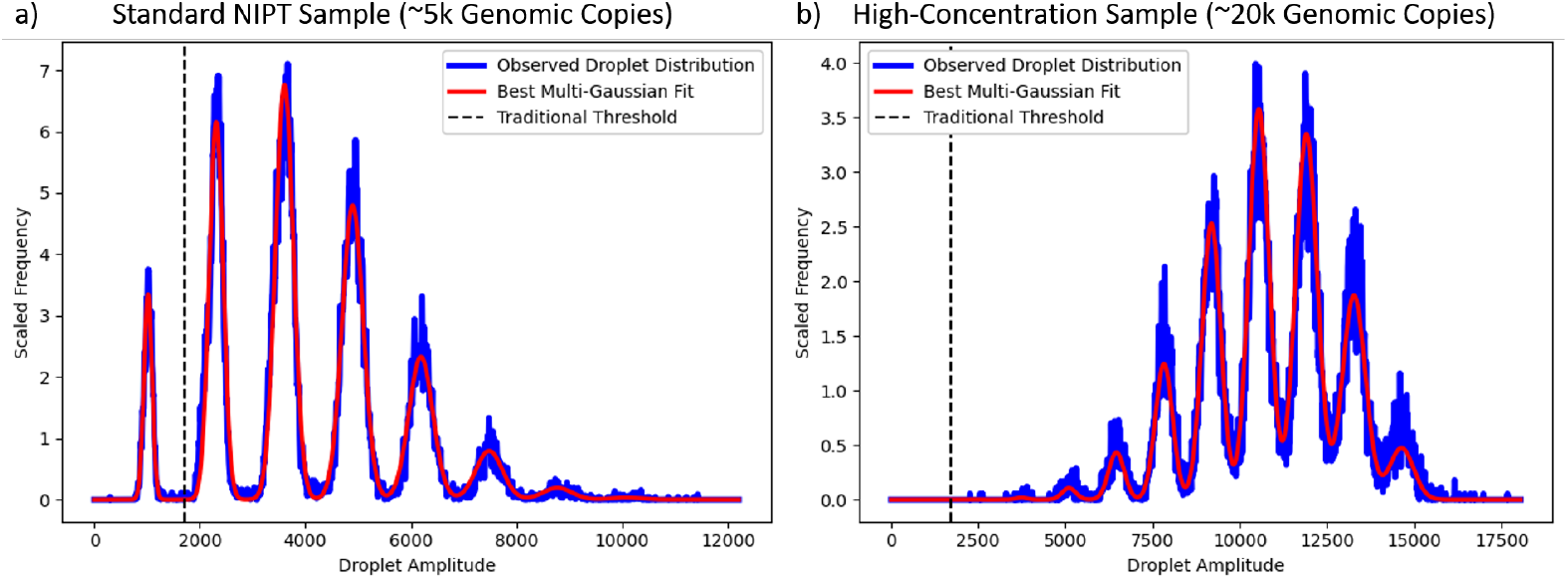
Multi-Gaussian Fitting. Rather than analyzing droplet amplitudes using a traditional positive/negative cutoff threshold (dashed black lines), using VPdPCR we fit the whole amplitude distribution to an extrapolated series of Gaussian functions (red lines). a) Our model matches the observed distribution well in a contrived cfDNA sample with significant peaks up to level 5. b) In a very high-concentration sample there is no 0-target peak, so traditional threshold analysis would fail completely. However, our multi-Gaussian fitting method is still able to perform an appropriate fit and thereby extract target concentration using VPdPCR analysis.

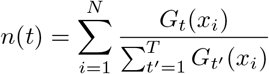

### Creating Virtual Partitions

Once *n*(*t*) has been determined for all bins we use it to calculate target copies. To do this we divide each of our *N* total partitions into *T* “virtual” partitions, of which *t* are positive and *T*−*t* are negative. This effectively transforms our *T*- target sample with *N* partitions into a 1-target sample with *T* N* partitions. We then count the negative virtual partitions and apply our formula from above to get target concentration:

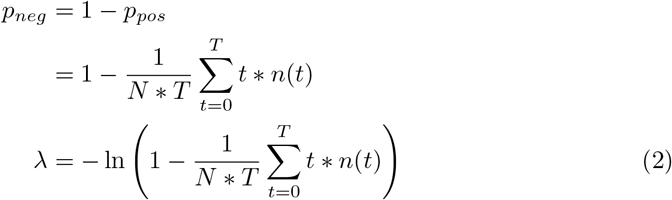

This formulation uses information from all bins 0 through *T* rather than just bin 0, allowing for accurate analysis at higher concentrations where few partitions are negative for all targets. As shown in Figure 6b this method even works when no droplets are present in the 0-target bin, a regime in which traditional threshold-based analysis breaks down completely.

### Theoretical Error Limits

Before experimentally evaluating the VPdPCR assay, we first determined theoretical optimum performance under different levels of multiplexing. For fetal trisomy testing the relevant analysis output is not the absolute number of copies of any one target but rather the ratio between total copies from one chromosome and total copies from another chromosome. The goal is to be able to consistently distinguish between a chromosome ratio of 1 (corresponding to a euploid mother and fetus) and a higher ratio corresponding to a euploid mother and fetus with a trisomy. Detailed statistical analysis by Dube et al [10] allows us to obtain 95% confidence intervals for calculated chromosome ratios given various true input ratios and partition counts. Figure 7a shows these confidence intervals for 20,000 partitions given simulated samples with a euploid fetus and a triploid fetus with a 5% fetal fraction in extracted cfDNA. At 7000 input genomic copies the intervals significantly overlap, indicating that a traditional assay with 1 target per chromosome cannot consistently identify fetal trisomy at a fetal fraction of 5%. If we increase the number of target regions per chromosome but maintain traditional threshold-based analysis we can effectively increase the input genomic copies without changing the number of partitions. However, as the graph shows, there is no input value for which the intervals are non-overlapping, indicating that no amount of multiplexing can make this task possible with threshold analysis.

**Fig 7.**
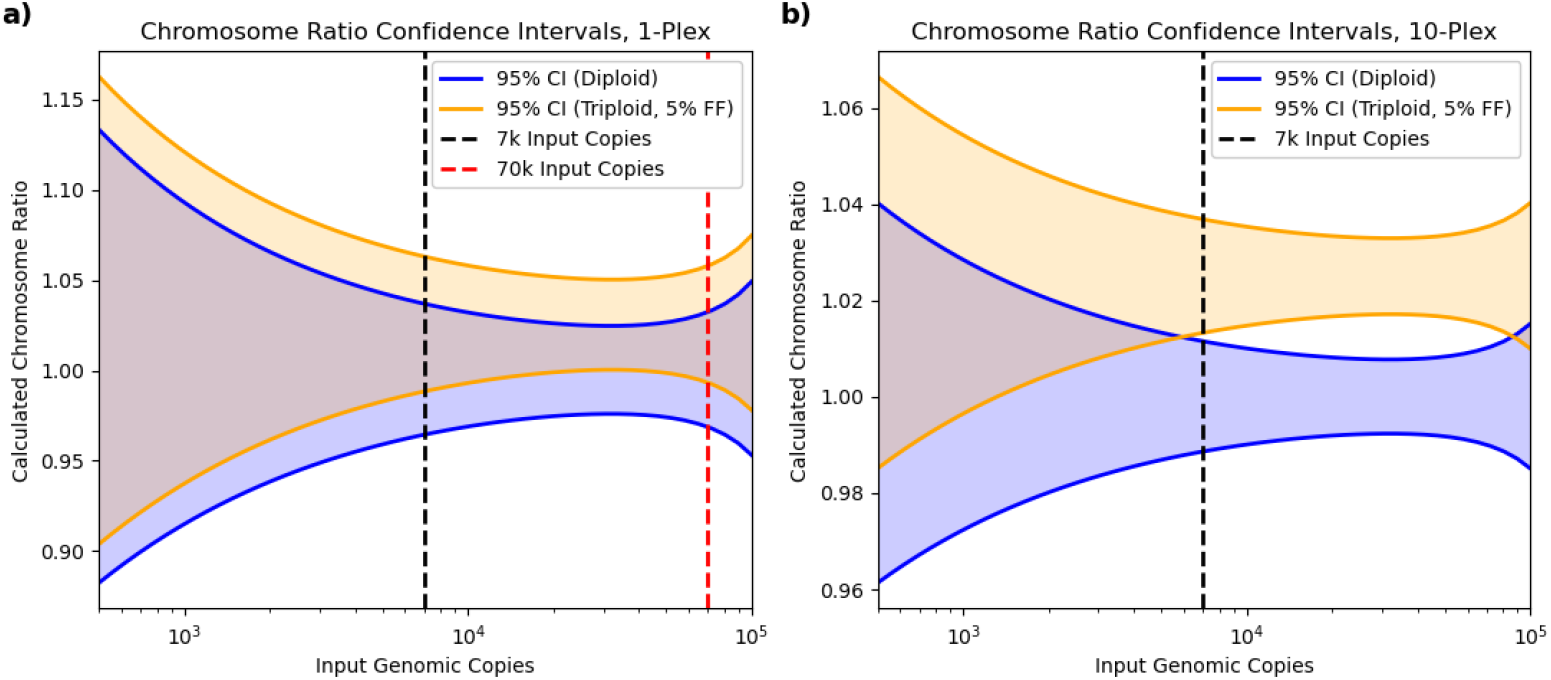
Theoretical Confidence Intervals. To consistently distinguish a diploid from a triploid fetus at a 5% fetal fraction in cfDNA, the two confidence intervals shown should be non-overlapping. a) With a singleplex assay on a machine with 20k physical partitions there is substantial overlap at 7k input copies. Multiplexing with traditional analysis is equivalent to a singpleplex assay with more input copies (red line), but no value of input copies produces non-overlapping intervals. b) Our 10-plex-per-chromosome VPdPCR assay expands the number of effective partitions, creating an input copy region with non-overlapping intervals and making it theoretically viable for fetal aneuploidy testing.

The conventional limitations of traditional threshold analysis changes significantly when we apply multi-Gaussian fitting and VPdPCR analysis to encompass all peaks in multiplexed reactions rather than simple positive/negative classification, which increases both the input copy number and the number of virtual partitions by a factor equal to the number of target regions per chromosome. Figure 7b-d shows the effect of this enhanced multiplexing on chromosome ratio confidence intervals. We found that 10 targets per chromosome is just enough to get theoretically consistent distinction between diploid and triploid samples at 5% fetal fraction and 7k input copies, so we chose that as the target of our assay design.

### Digital PCR Assay Design

Multiple TaqMan^®^ PCR assays were designed to amplify conserved regions of Chromosomes 18 and 21 using the Primer3 command line tool [30, 31] with GNU Parallel [32] to process designs efficiently on multiple computer cores. We also used primer3 to calculate the binding energies of all pairwise dimers ([monovalent cation] = 50mM, [divalent cation] = 2.5mM, [dNTP] = 0.8 mM, temperature = 60°C), and assays were removed to eliminate favorable oligo-oligo interactions until we reached 10 assays per chromosome. The selected primers and probes were ordered from Integrated DNA Technologies, Inc. (Coralville, IA). Chromosome 18 and 21 TaqMan probes were labeled to be detected in dye channel 1 and channel 2 respectively, and both target chromosomes’ TaqMan probes were double quenched with ZEN™ quenchers.. A 20-plex oligo mix was prepared with all the primers at equal concentration and probes at a significantly lower concentrations. The assay is in development and has not been officially released or approved by the U.S. Food & Drug Administration.

### Sample Preparation

Cell line DNA stocks from the Coriell Institute for Medical Research (NA04965 (Trisomy 21), NA12878 (euploid), and NA15453 (euploid)) were sheared with a Covaris^®^ E220 Focused-ultrasonicator (SKU500239) in order to have a mean length of 150 base pairs, simulating the short fragments found in cfDNA. Sheared DNA was processed through the standard singleplex ddPCR workflow using a chromosome 18 target to calculate concentration. To simulate presence of Trisomy 21 stocks were diluted in 1x, low EDTA TE (Gbiosciences 786-151) to a total of 3500 copies/5 *µ*L and 7000 copies/5 *µ*L at 5%, 10% and 20% NA04965 in NA12878. To simulate euploid samples the euploid cell lines were diluted to a total of 3500 copies and 7000 copies/5*µ*l at 0%, 5% and 100% NA15453 in NA12878.

### Digital PCR Methods

PCR reactions were set up using the following volumes: 10 *µ*L 2x ddPCR Supermix for probes (no dUTP) (BioRad Laboratories^©^ 186-3024), 5 *µ*L of 20plex oligo mix, and 5 *µ*L of sample. 15 *µ*L of PCR mix was added to each well of 96 well ddPCR plate (BioRad Laboratories 12001925) followed by 5 *µ*L of each sample. Plates were sealed using pierceable foil heat seal (BioRad Laboratories 1814040) and the PX1 plate sealer (BioRad Laboratories 1814000). Plates were vortexed, spun down, and run on the Automated Droplet Generator (BioRad Laboratories 1864101) using Automated Droplet Generation Oil for Probes (BioRad Laboratories 1864110) and the DG32^™^ Automated Droplet Generator Cartridge (BioRad Laboratories 1864108). After droplet generation was completed, thermocycling was performed on the C1000 Touch with the 96 deepwell module (BioRad Laboratories 1840197). Thermocycling was performed as follows: 1. Enzyme activation (95°C for 10 minutes), 2. 45 Cycles consisting of denaturation (95°C for 20 seconds) followed by combined annealing/extension (58°C for 2 min), 3. Enzyme deactivation (98°C for 10 minutes), and 4. A 4°C infinite hold. Signal detection was performed on the QX200 Droplet Reader (BioRad Laboratories 1864001). Experiment was set to ABS, Target 1 was set to Ch1 Unknown, Target2 was set to Ch2 Unknown and Supermix was set to ddPCR Supermix for probes (no dUTP). Wells were read in columns. Data was exported using the BioRad Laboratories QuantaSoft Version 1.7.4.0917 Software and analyzed using the Python Programming Language version 3.7 (Python Software Foundation, https://www.python.org/).

## Conclusion

Digital PCR enables best-in-class rapid quantitative percision for monitoring genomic fragments. However, dPCR on its own, is insufficient for some clinical diagnostic applications, including non-invasive prenatal testing. VPdPCR method not only enhances single-well dPCR multiplexing by a factor of 10 in this demonstration, but also enables dPCR platforms to overcome fundamental limitations to precision by decreasing both sampling and partition error. With newer multi-channel digital platforms, VPdPCR could enable a complete aneuploidy panel for chromosomal abnormalities (Chr21, Chr18, Chr13, X, Y) in a single well. We believe the enhanced precision of VPdPCR could also be useful in a variety of other diagnostic settings, such as detecting copy number variation of crucial genes to perform liquid biopsies or analyzing low abundance mRNA expression. This range of applications has the potential to make VPdPCR a standard of practice for precision molecular diagnostics.

## Supporting information

**Table S1.**
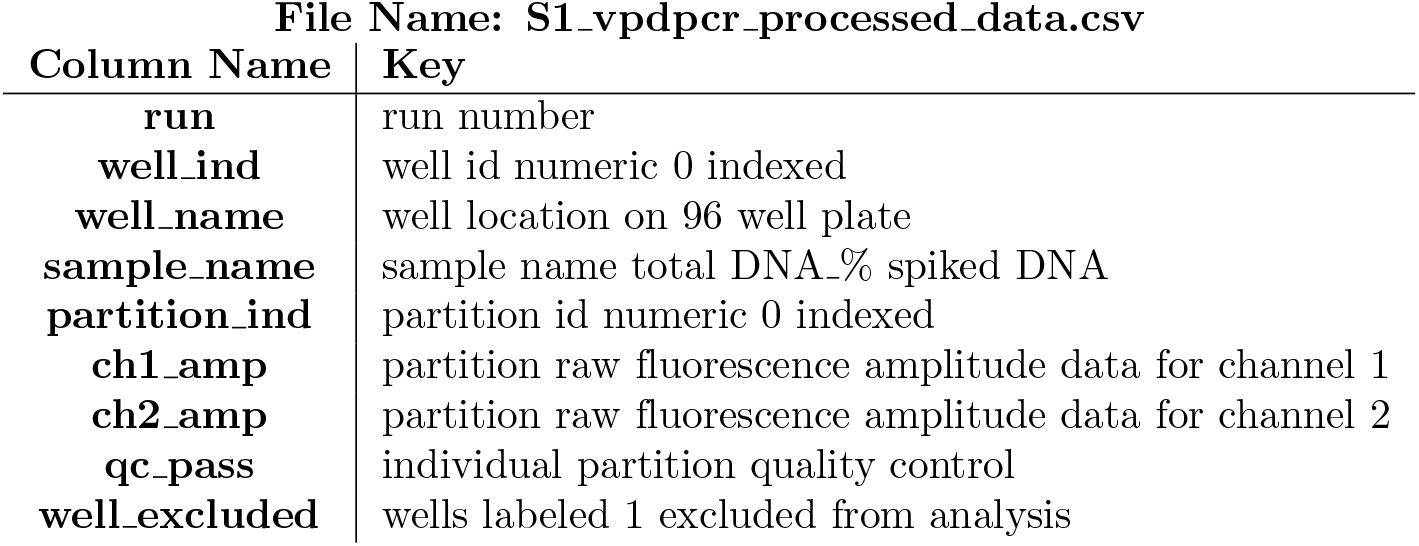
Digital 20plex Processed Data Table Key

## Acknowledgments

The authors would like to thank Jeff Gole and Mimi Wang of ChromaCode, Inc. for making software available to select conserved genomic regions for primer design, Sheila Rosenburg of ChromaCode, Inc for critique and lively statistical discussions and Molly Smith of ChromaCode, Inc for procuring the sheared DNA from UCSD. The following cell lines/DNA samples were obtained from the NIGMS Human Genetic Cell Repository at the Coriell Institute for Medical Research: NA12878DNA, NANA15453DNA and NA04965DNA. Covaris DNA Shearing was conducted at the IGM Genomics Center, University of California, San Diego, La Jolla, CA.

